# Lepidoptera Diversity, Richness, and Distribution in Semi-Urban Farmland and other Habitats around Lumbini, Rupandehi

**DOI:** 10.1101/2025.01.26.634957

**Authors:** Celeus Baral, Hem Sagar Baral, Carol Inskipp, Rajana Maharjan

**Affiliations:** NAMI College, University of Northampton, Kathmandu Nepal; Himalayan Nature, Kathmandu, Nepal; School of Veterinary, Environmental and Agriculture Sciences Charles Sturt University, Australia

**Keywords:** Habitat, Lepidoptera, Species Diversity, Semi-urban Farmland, Environmental Factors, Distribution Patterns

## Abstract

This study comprehensively examined Lepidoptera diversity in the semi-urban agricultural landscapes of Lumbini, Nepal, documenting 30 moth species from six families and 39 butterfly species from five families over 14 days. Field methods included moth traps with 50 Watts LED lights inside a box with egg cartons for moth trapping and butterfly transect surveys across various habitats—grasslands, shrublands, and agricultural areas for recording butterfly species. The Erebidae family dominated moth populations, while Nymphalidae led butterfly diversity, showing resilience in disturbed habitats influenced by environmental factors such as weather patterns, lunar phases, and habitat characteristics. Statistical analyses using Simpson’s Diversity Index, Shannon-Wiener Diversity Index, Pielou’s Evenness Index, and Margalef’s Richness Index revealed that shrubland and agricultural habitats supported the highest species richness and evenness, while grasslands hosted fewer species. The study highlighted the importance of agricultural and shrubland ecosystems for biodiversity, emphasizing ongoing monitoring to understand how Lepidoptera populations respond to environmental changes, offering essential insights for future conservation efforts.

## Introduction

Lepidoptera, comprising moths and butterflies, is one of the largest and easily recognizable insect orders, with around 160,000 species globally [41]. Characterized by their scales and proboscis, they undergo complete metamorphosis in four stages: egg, larva (caterpillar), pupa, and adult [17]. Moths, accounting for the majority of species, belong to 30 superfamilies, with families like Noctuidae (about 35,000 species) and Geometridae (around 21,000 species) being the largest [37]. On the other hand, butterflies are divided into 46 superfamilies, with Nymphalidae and Lycaenidae (around 6000 species) being the most prominent [23]. Moths are mainly nocturnal, butterflies are diurnal and most species are often more vibrant than moths. Both groups are vital pollinators, particularly moths for night-blooming flowers, contributing to plant reproduction and ecosystem stability [42]. Additionally, moths influence plant population control and nutrient cycling. Butterflies, known for their beauty, assist in long-distance pollen distribution, boosting plant genetic diversity [18]. Due to their sensitivity to environmental changes, Lepidoptera are often used as ecological indicators [12]. Conservation efforts often prioritize Lepidoptera due to their public appeal, but focusing on diversity rather than specific taxa can benefit broader ecological communities [36]. Additionally, these insects are critical for ecosystem health, evolutionary biology, and pest management, highlighting their ecological importance.

Butterflies play an important part in pollination by dispersing pollen while feeding on nectar. Tiny scales on their bodies collect pollen from one bloom and transport it to another, facilitating plant reproduction. Butterflies and moths, both are closely linked to flowering plants (angiosperms), which are crucial for their reproduction. Butterflies’ unusual proboscises allow them to consume nectar from various flowers, fostering genetic diversity in plants by dispersing pollen across great distances [24]. While some species have specific flower preferences, many are adaptable, selecting plants based on flower color, nectar content, and structure [60].

Approximately 87.5% of blooming plants rely on animal pollinators, with nocturnal moths being especially important yet sometimes underestimated [69]. Moths from the families Sphingidae, Noctuidae, and Geometridae are important evening pollinators in tropical rainforests and temperate woods [27]. Their involvement in pollination promotes interpopulation gene flow, long-distance pollen dispersal, and efficient pollen transfer. Unlike butterflies, which are active during the day and favor vividly colored flowers, moths’ nighttime activities highlight their critical role in pollination [33]. Understanding the pollination roles of moths and butterflies is critical to preserving these ecological linkages and devising successful conservation measures.

Lepidoptera face increasing vulnerability due to climate change, but evidence documenting these impacts in Nepal is scarce. Globally, climate change threatens many Lepidoptera species; Wilson and Maclean [67] predict that 10% could become vulnerable by the century’s end due to specialized habitat needs and fragmented ranges. While some species might expand northward, those in lower latitudes or mountainous areas may face habitat loss and limited migration options. Fox [14] reports a 31% decline in moth populations in Britain over 35 years, exacerbated by habitat degradation from intensive farming and chemical use. Wagner [64] notes similar declines in the northeastern U.S., driven by habitat loss, increased predation, and potentially climate change. The lunar cycle also influences moth activity, reducing attraction to artificial light during full moons [68]. Moths in India, particularly Lasiocampidae, have also declined due to habitat loss, diseases, and parasites [1;19]. In contrast, butterflies, while also declining, benefit from extensive documentation. Rödder *et al.* [45] observe that European butterflies are moving to higher elevations due to rising temperatures, while Crossley *et al.* [10] find declines in North American butterflies linked to increased dryness and heat. Kwon et al. [31] highlight habitat changes from reforestation and urbanization in South Korea as more impactful than climate change. In Nepal, Gaudel *et al.* [16] report a decline in butterfly populations along Mahendra Highway due to roadkill. Addressing these issues requires extensive research to understand Lepidoptera’s ecological roles and inform conservation strategies.

Nepal’s diverse habitats, spanning tropical Terai plains to the alpine regions of the Himalayas, support a wide range of Lepidoptera [9]. In the tropical and subtropical zones, especially in the Terai, dense forests provide ample feeding and breeding opportunities for many species. The mid-hills shrublands and grasslands, characterized by various grasses and low-lying plants, host an abundance of butterflies and moths. Lepidoptera thrives on crops and surrounding vegetation in agricultural areas, while temperate forests in the mid-hills support species adapted to cooler climates. Alpine meadows in the Himalayas are home to specialized species that survive in high-altitude conditions [53]. Even urban gardens offer crucial habitats, making Nepal an ideal region for studying Lepidoptera diversity across varied ecosystems. Around 663 butterfly species have been identified in Nepal [47] and 6,000 macro moth species have been recorded [53]. The diversity of habitats found in Nepal’s climatic and geographical zones reflects the country’s ecological complexity and richness in Lepidoptera. Each ecosystem supports distinct types of butterflies and moths, making Nepal a perfect place for researching these critical ecological markers.

Although Nepal has a rich biodiversity and has been recognized for successful conservation and habitat management efforts, such as the recent increase in tiger populations in the lowlands [13], smaller species such as butterflies and moths are frequently overlooked and not prioritized. Nepal’s research has focused mostly on larger animals, leaving a considerable gap in insect studies [25]. While few species-specific studies have been on Lepidoptera of Nepal [39; 28] comprehensive diversity studies remain rare, particularly for moths. The diversity and richness of moths in different regions of Nepal are still barely studied, emphasizing the necessity of comprehending the functions of these essential ecological indicators across a range of habitats, elevations, and climates.

Lumbini’s rich natural history and diverse ecosystems—comprising farmlands, wetlands, forests, and shrublands—are vital for sustaining various forms of life [3]. Studying the diversity, richness, abundance, and distribution of lepidopterans, serving as important ecological indicators, is essential for informing conservation policies and assessing the region’s environmental health [60]. Given Nepal’s varied terrain, this area has significant potential for further research. Enhancing the understanding of moth and butterfly status, roles, and ecological services in Nepal will contribute valuable insights to biodiversity conservation efforts.

The study aims to accomplish three key objectives: a) Assess species diversity and richness of Lepidoptera in Lumbini Sanskritik Municipality of Rupandehi District, focusing on the variety of butterfly populations across habitats such as agricultural fields, forests, wetlands, grasslands, and shrublands, b) investigate how varied ecosystems influence butterfly species and family distributions, analyzing the impact of habitat types—such as agricultural lands, shrublands, forests, and wetlands—on butterfly distribution and abundance, and c) examine the relationship between moth diversity, richness, and evenness with environmental factors like weather conditions and moon phases. The findings will offer insights into the ecological preferences of these insects, enhancing understanding of their role in the ecosystem.

## Materials and Method

### Study Area

The study was carried out at Lumbini, in the Rupandehi district of Lumbini Sanskritik Municipality, Lumbini Province. Lumbini, the birthplace of Lord Buddha, is a site of immense historical and religious importance, housing archaeological artifacts from the third century BC. Key structures include the Shakya Tank, Maya Devi Temple, and the Ashoka pillar with Brahmi inscriptions reflecting the region’s rich history [63]. The study area was centred at coordinates 27°29’24" N and 83°17’47" E, approximately 100 meters above sea level. The total area of research covered was approximately 6,771.84 square meters.

**Figure 1:**
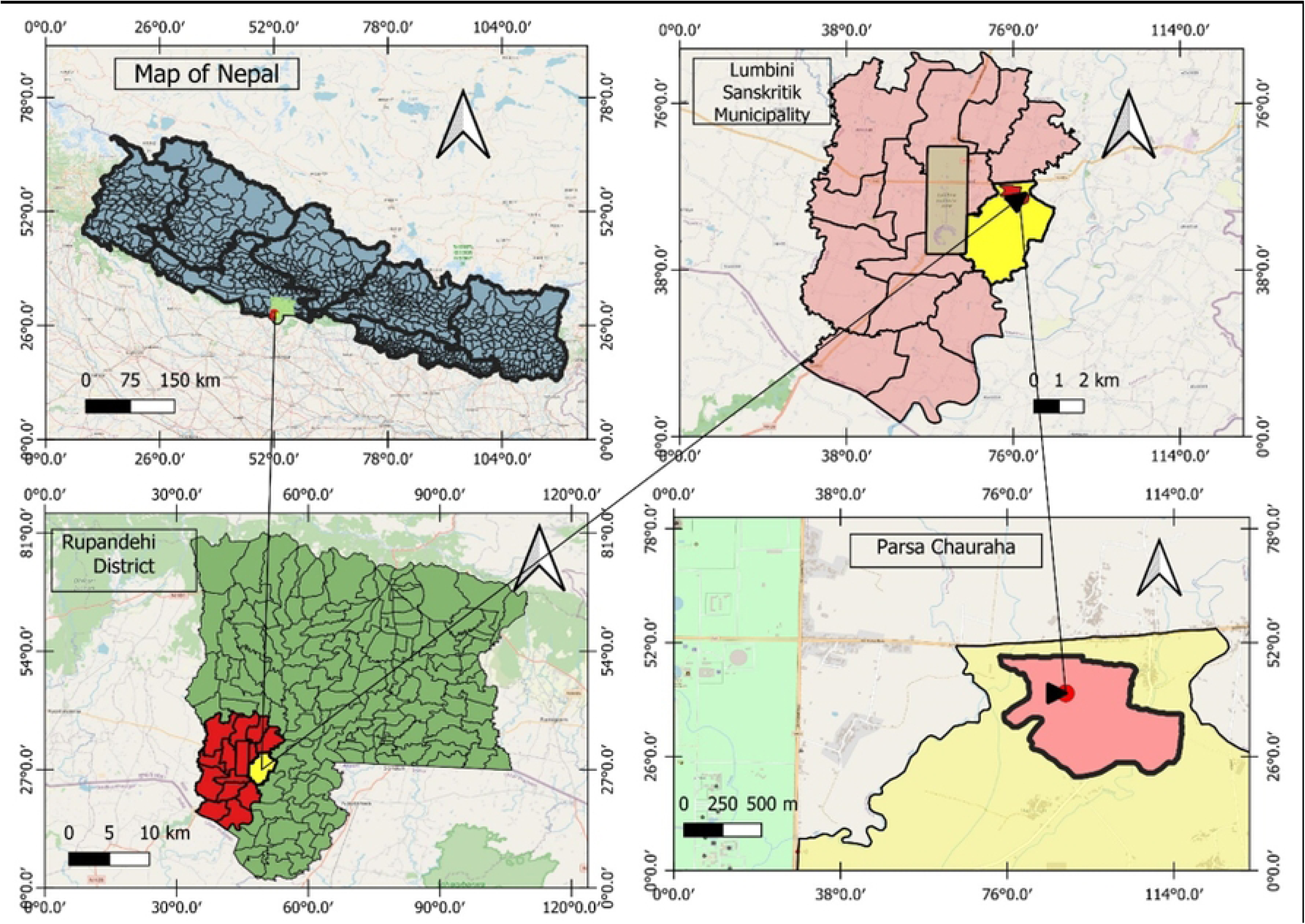
The study area map illustrating Lumbini Sanskritik Municipality, Rupandehi District.

**Figure 2:**
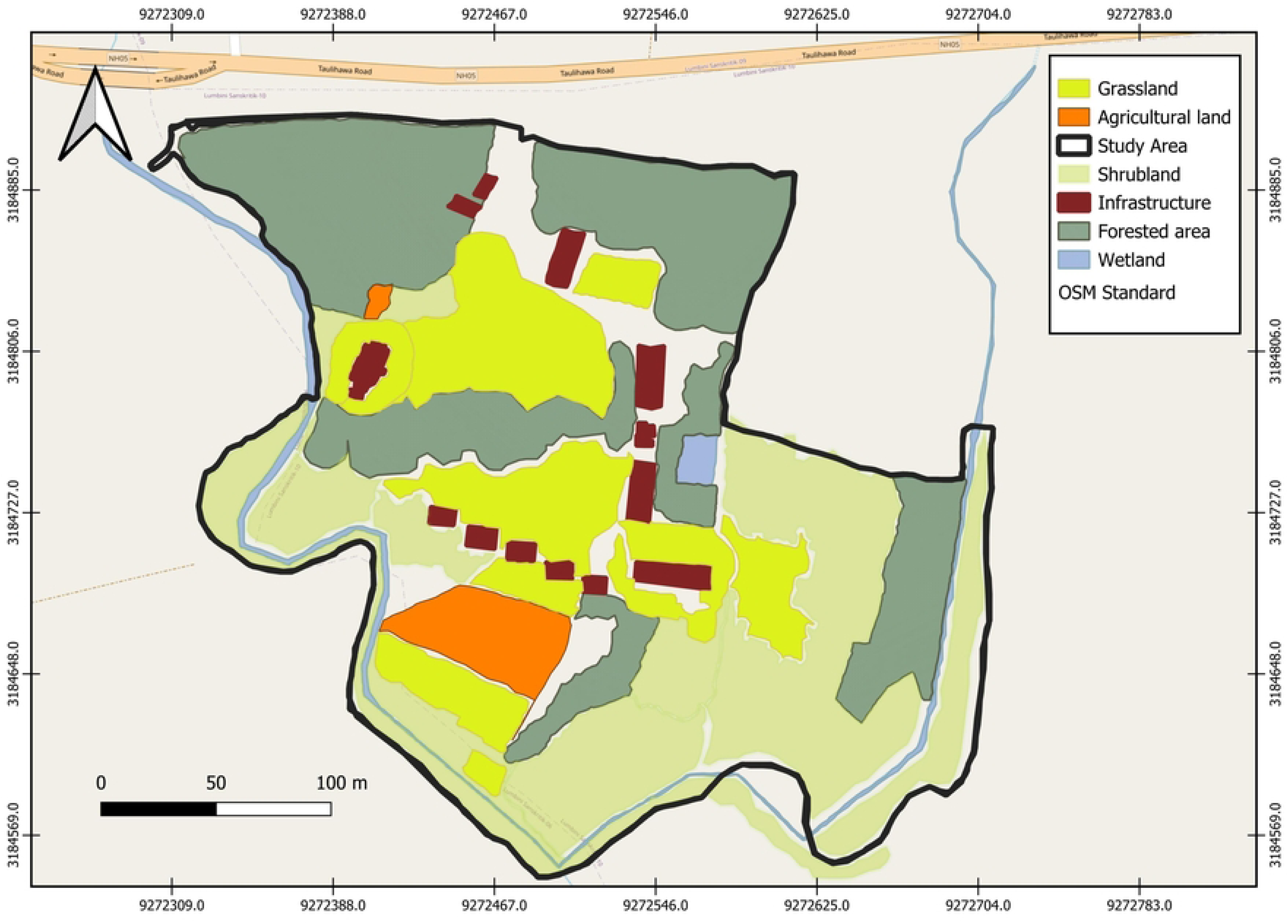
A polygon map of the habitat distribution across the study area.

### Climate and Vegetation

Lumbini experiences a tropical to subtropical climate with distinct seasons, including long, hot summers that often exceed 40 °C, and a prolonged monsoon from June to September while winters can drop as low as 9 °C, offering a cooler period essential for agriculture. In recent years, temperatures have risen by about 4 °C due to global warming [3], posing threats to biodiversity and potentially Lepidoptera species. Higher temperatures affect water availability and growing seasons [34].

Lumbini encompasses diverse habitats, including open water bodies (10 ha), forest plantations (270 ha), and grasslands (400 ha) [3]. Lumbini Development Trust, a local authority in the area, has planted mixed types of trees, native and imported [3]. The area supports 72 vascular plant species, mainly from the Asteraceae, Poaceae, and Fabaceae families, with Fabaceae being culturally significant due to its association with Lord Buddha [6]. However, human activities and alien species introductions have led to species decline [51]. Key nectar-rich plants include *Vernonia cinerea* and *Helianthus annuus*, while Asteraceae, Lamiaceae, and Moraceae provide essential resources for butterflies [51]. Lumbini is also home to 421 bird species, including the globally threatened Sarus Crane [3], as well as mammals such as Nilgai *Boselaphus tragocamelus*, Wild Boar *Sus scrofa* and Indian Grey Mongoose *Urva edwardsii*. The Lumbini farmlands and Jagadishpur Reservoir are designated as Important Bird and Biodiversity Areas of Nepal [4].

### Data Collection

The study was carried out from 19 February to 5 March 2024 with a moth trap placed in a semi-natural cultivated lemon farm/shrubland. The moth trap of 50W LED light was fitted inside a cardboard structure and was set up to capture moths at a semi-urban rural site. This site was chosen for its plant variety and proximity to populated areas—factors influencing moth activity. The trap included egg cartons and two glass panes, ensuring trapped moths stayed in the box. The light was turned on the whole night and was checked every morning at 6.30 AM. Moths collected were photographed under secured conditions to ensure their safety and facilitate accurate identification.

Butterfly diversity data was gathered through field surveys and direct observation. On the first day, monitoring occurred from sunrise to sunset, recording butterfly movements and activities hourly to capture temporal variations. Subsequent days focused on peak activity hours from 12:00 PM to 3:00 PM. Surveys covered a 500-meter transect through grasslands, shrublands, wetlands, and forests, with species identified and photographed when possible. Photographs aided identification of the species during analysis.

Species identification for both moths and butterflies used digital resources like iNaturalist and field guides by Smith [53, 54] and Gurung *et al*. [20], with genus-level classification applied when species-level identification was challenging. This method was thoroughly followed for at least 14 days to build a robust dataset. Habitat descriptions and dominant plant species were crucial for interpreting both moth trap and butterfly transect results.

**Figure 3a.** The moth trap and its setting.

**Figure 3b.** The 50-watt LED light lit up during nighttime.

**Figure 4a.** Study conducted based on observation.

**Figure 4b.** *Ideopsis vulgaris* observed.

The observed Lepidoptera species were classified into four abundance categories: Very Rare (VR) for 1-2 individuals, Rare (R) for 3-10, Fairly Common (FC) for 11-30, and Common (C) for more than 30 individuals. This systematic classification helped assess species prevalence within the study area [52].

### Data Analysis

#### 3.5.1. Simpson’s Diversity Index (D)

The data was examined using a combination of Microsoft Excel 2013 and R Studio for statistical accuracy. Simpson’s Diversity Index (D) was used to quantify species diversity, determined by the formula:

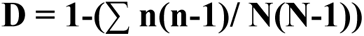

Where,

n = Number of individuals of a species

N= The total number of individuals of all species

D= Simpson’s Diversity Index

This formula measures the likelihood that two randomly selected individuals belong to different species. Index values range from 0 to 1, with higher values indicating higher diversity [55].

#### 3.5.2. Shannon-Weiner Diversity Index (H’)

The Shannon-Wiener Diversity Index (H’) was also used to assess both species richness and evenness:

The formula for the Shannon-Weiner Diversity Index (H’) is:

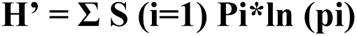

Where,

S= Total number of species

pi = the proportion of individuals of species iii relative to the total number of individuals (i.e., pi=ni\N, where ni is the number of individuals of species i, and N is the total number of individuals)

ln(pi) represents the natural logarithm of *pi*. The Shannon-Wiener index accounts for both species richness and evenness, with higher H′ values indicating greater diversity [7].

#### 3.5.3. Pielou’s Evenness Index (J’)

The evenness was calculated to reveal the relative abundance of species distributed using Pielou’s Evenness Index (Equitability) (J’).

The formula for the Pielou’s evenness index (J’) is:

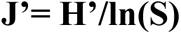

Where,

H′ = Shannon-Wiener diversity index,

S = the total number of species, and

ln(S) = natural logarithm of S.

Pielou’s Evenness Index assesses how equally species are distributed among species in a group. It has a range of 0 to 1, with values closer to 1 suggesting more evenly distributed species [21].

#### 3.5.4. Margalef’s Richness Index (Dmg)

Margalef’s Richness Index was calculated to measure species richness by adjusting for the total number of individuals obtained, giving an accurate comparison across several habitats.

The formula for the Magalef’s Richness Index (Dmg) is:

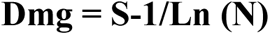

Where,

S = Total number of species.

N = Total number of persons.

ln(N) = The natural logarithm of the total number of species.

Margalef’s index quantifies species richness, with higher values reflecting greater richness relative to sample size [15]. R-studio was used to create Rank-abundance (Whittaker) plots for moths and butterflies, while Q-GIS was used to generate the maps.

## Results

### Moths

A total of 30 moth species from six families and 26 different genera was observed. The Erebidae family was the most prevalent, representing 44 individuals from various genera, followed by Noctuidae with 29 individuals. Other families, including Geometridae, Crambidae, Pyralidae, and Spilomelinae, had fewer abundances, contributing 5, 4, 3, and 1 individual, respectively. Notably, the Tiger Moth (*Spilosoma strigatula*) of the Erebidae family and the Asiatic Pink Stem Borer (*Sesamia inferens*) of the Noctuidae family were both categorized as FC, with 25 and 18 individuals documented, respectively.

Erebidae represented for 43.33% of the total, dominating the habitat. In comparison to Erebidae, Noctuidae appeared second, with a 26.67% prevalence, indicating a notable but less dominance (Figure 5). A reasonable distribution of these families may be seen in the 10% occupation from Geometridae and Crambidae, respectively. Spilomelinae was the smallest group, accounting for 3.33%, while Pyralidae comprised 6.67%. An illustration of the ecological niches or environmental conditions accompanied the distribution of each family’s abundance.

**Figure 5:**
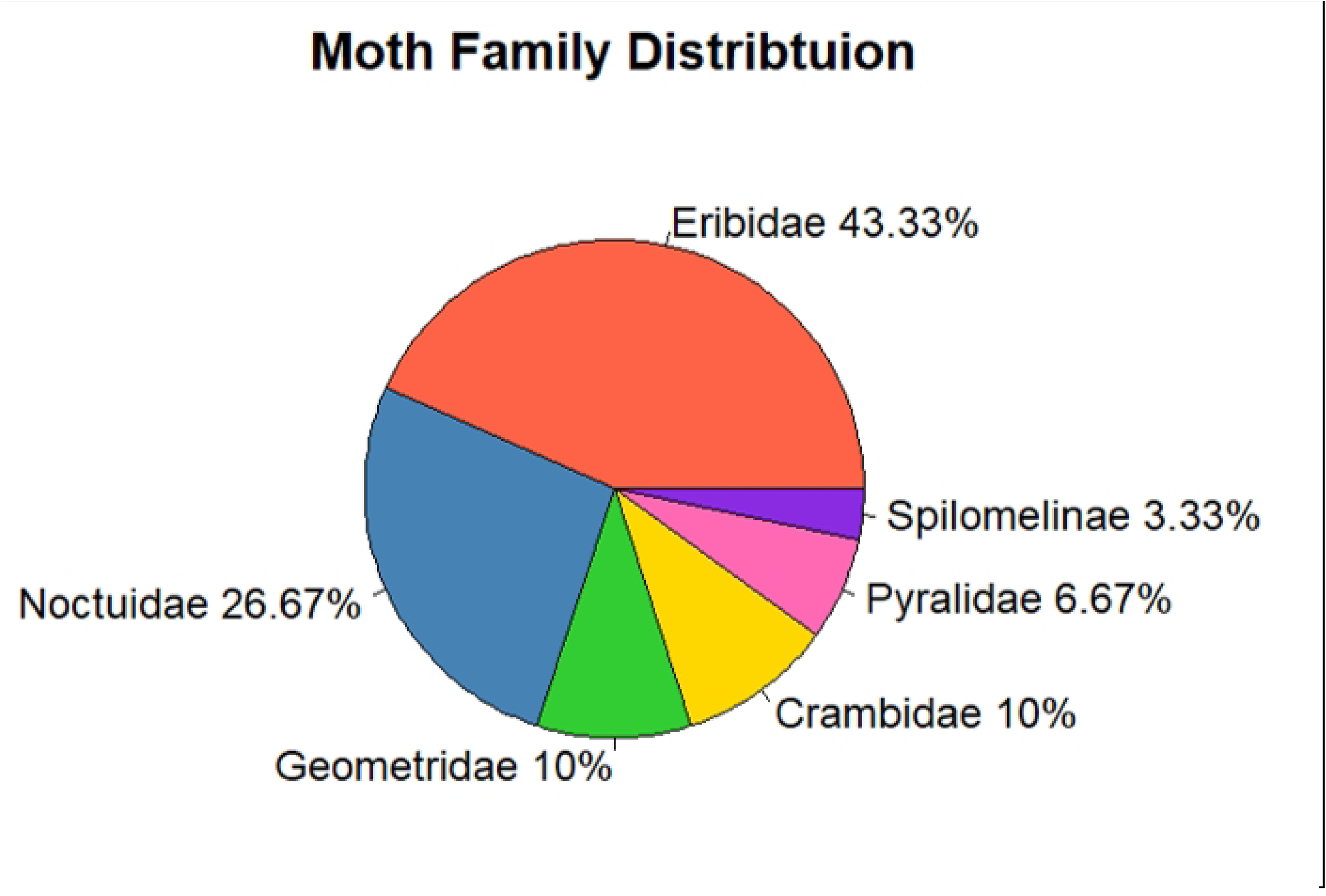
The distribution of moths by the families in percentage.

A clear pattern of uneven species distribution among moths was found (Figure 6). Spilosoma strigatula and Sesamia inferens appeared dominant, with relative abundances of approximately 0.3 and 0.23, respectively. These accounted for a significant portion of the population, demonstrating their prevalence in the moth community. Following these dominant species, there is a sharp drop in abundance, with *Asura-Miltochrista* group sps, ranked third. The mid-ranked species, such as *Dasychira* and *Parapoynx diminutalis*, exhibited varying levels of abundance, ranging from 0.034 to 0.023. This variation suggests that while these species are not as dominant, they occupy intermediate ecological roles, contributing to the diversity but still maintaining relatively low abundances. Species such as *Creatonotos gangis-interrupta* and *Eressa confinis* highlighting the presence of rare species within the community. This flattening reflects a long tail of rare species that contribute little to the overall population but add to the species richness. Overall, the plot highlighted a highly uneven distribution within the moth community, with a few species dominating while the majority being rare.

**Figure 6:**
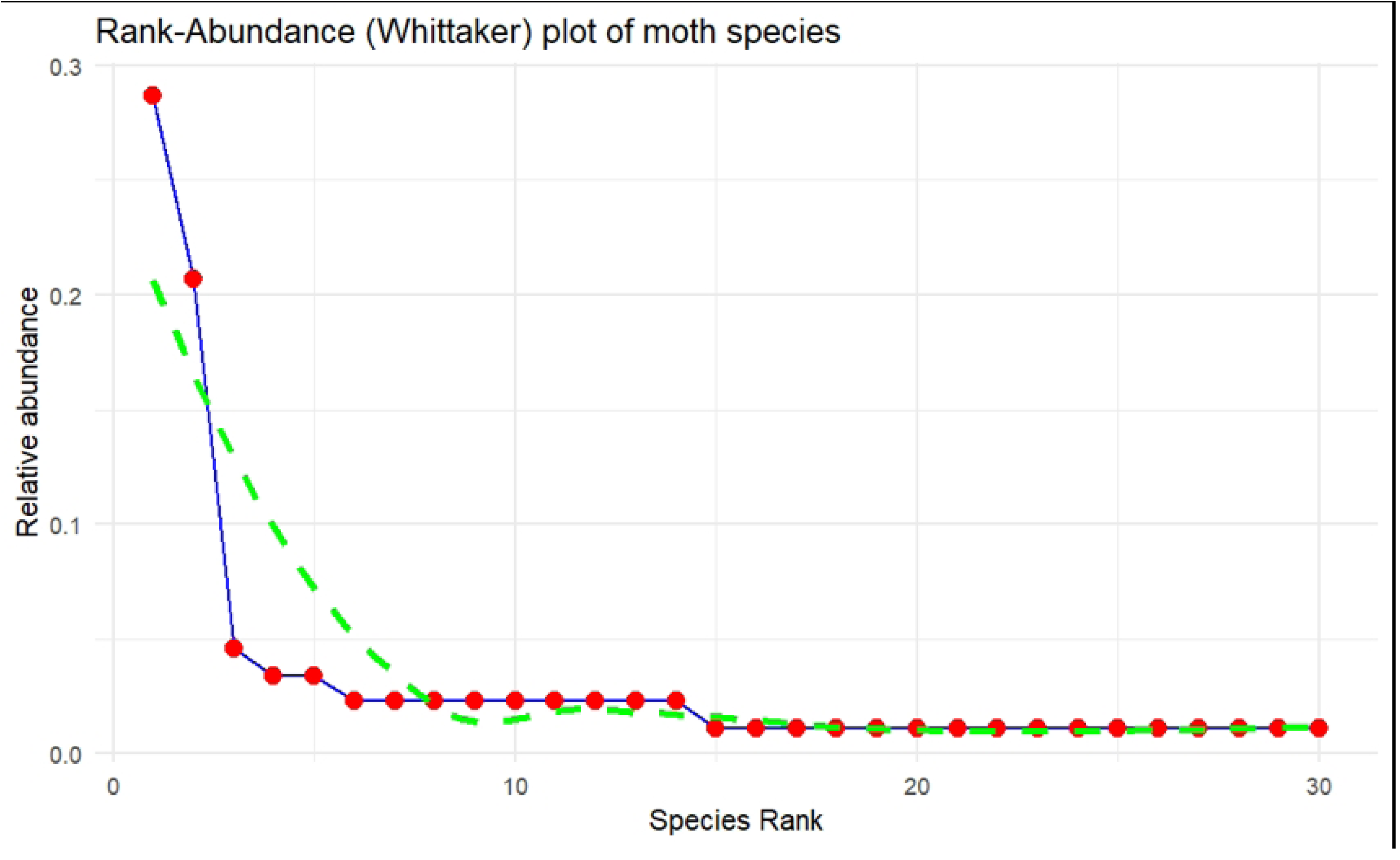
Rank-abundance plot of moth species observed.

Table 1 represented Eribidae, contributing 13 species, having the highest diversity (H’ = 1.73), dominance (D = 0.67), and species richness (Dmg = 3.15). Noctuidae, with moderate richness (Dmg = 2.08) and diversity (H’ = 1.36), has lower dominance (D = 0.59) and evenness (J’ = 0.66). Crambidae and Geometridae exhibit similar diversity and dominance, though Crambidae is more even (J’ = 0.95) and Geometridae slightly richer (Dmg = 1.24). Pyralidae has the lowest richness (Dmg = 0.91) and diversity (H’ = 0.64), but high evenness (J’ = 0.92). Spilomelinae, represented by a single species, lacks diversity indices. The overall community diversity (H’) is 2.66, with dominance (D = 0.86) and moderate evenness (J’ = 0.78).

**Table 1:** The Shannon-Weiner index (H’); Simpson’s Diversity Index (D); Margalef’s Richness Index (Dmg); Pielou’s Evenness (J’) and Species Richness (S) of all of the 6 families and their overall total.

| Family | H' | D | Dmg | J' | S |
| --- | --- | --- | --- | --- | --- |
| Eribidae | 1.73 | 0.67 | 3.15 | 0.67 | 13 |
| Noctuidae | 1.36 | 0.59 | 2.08 | 0.66 | 8 |
| Crambidae | 1.04 | 0.63 | 1.44 | 0.95 | 3 |
| Pyralidae | 0.64 | 0.44 | 0.91 | 0.92 | 2 |
| Geometridae | 1.05 | 0.64 | 1.24 | 0.96 | 3 |
| Spilomelinae | 0 | 0 | 0 | 0 | 1 |
| Overall Total | 2.66 | 0.86 | 6.49 | 0.78 | 30 |

### Butterflies

A total of 39 butterfly species from five families and 31 different genera was observed. The Pieridae family emerged as the most abundant, with 969 individuals spanning multiple genera, followed by Lycaenidae, which accounted for 720 individuals (Appendix 3). In contrast, other families such as Papilionidae, Nymphalidae, and Hesperiidae showed significantly lower abundances, contributing 227, 199, and 3 individuals respectively. Notably, the Large Cabbage White (*Pieris canidia*) from the Pieridae family and the Lesser Grass Blue (*Zizina otis*) from Lycaenidae stood out, both classified as “Common” due to their high counts of 578 and 536 individuals, respectively.

Nymphalidae was the most dominant butterfly family as per the diversity, comprising 41.03% of the total population, indicating its significant presence across the various habitats such as grasslands, shrublands, agricultural field, wetland and forest (Figure 7). Pieridae represented 25.64% of the population, reflecting its relatively high abundance. Lycaenidae accounted for 15.38%, while Papilionidae and Hesperiidae were less prevalent, making up 10.26% and 7.69%, respectively. This distribution suggested that certain families, such as Nymphalidae and Pieridae, were better adapted or more abundant in these habitats, while others maintained a smaller but notable presence.

**Figure 7:**
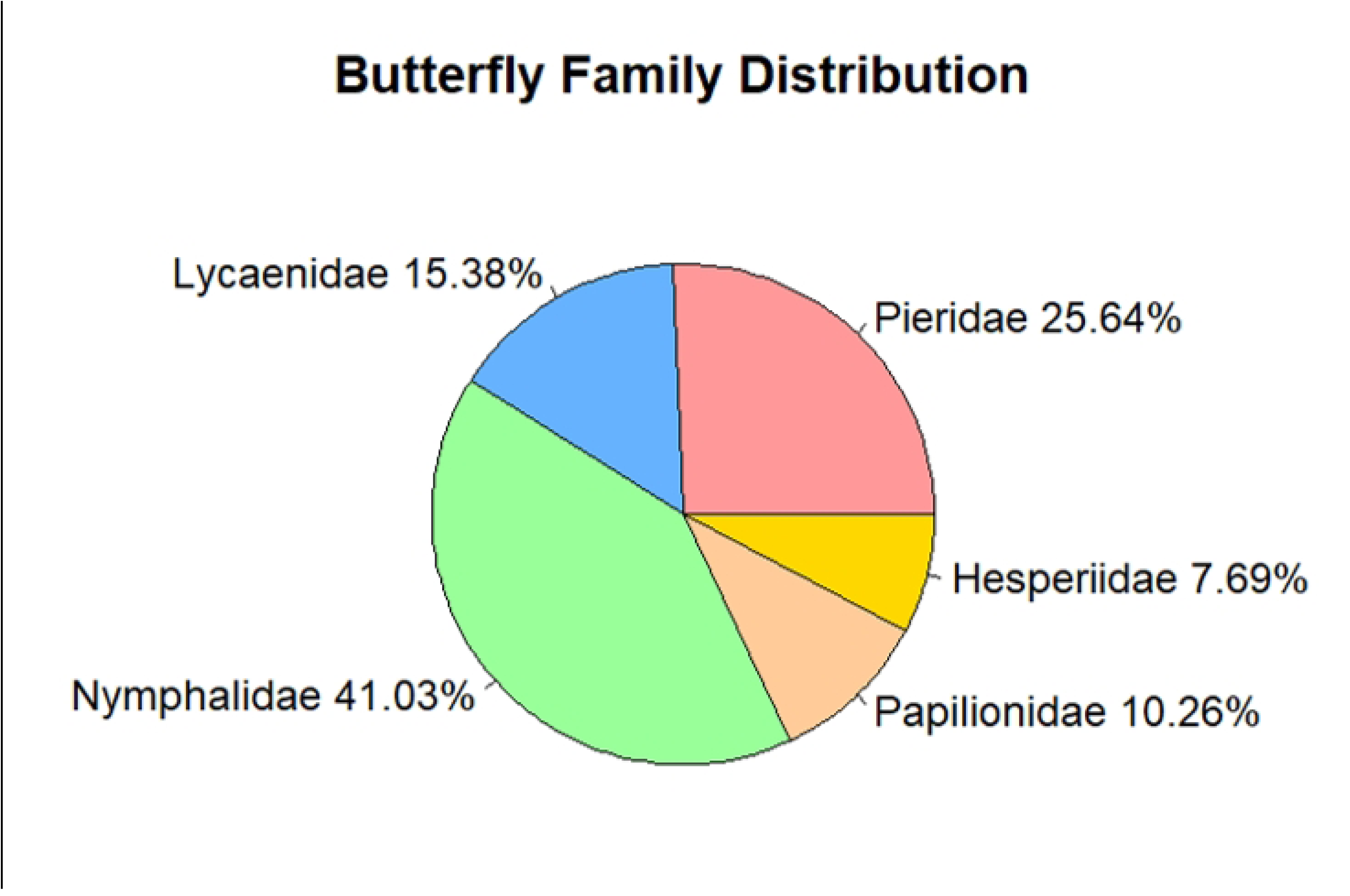
The distribution of the butterflies by the families in percentage.

Species such as *Pieris canidia*, *Zizina Otis* and *Papilio polytes* had higher abundances compared to others (Figure 8). After the top five dominant species, abundance declined steeply for others indicating that only a few species were numerically dominant such as the *Danaus genutia* and *Gonepteryx rhamni*. Species ranked beyond 10 contributed very little to the overall population such as the *Hypolimnas bolina* and *Cepora nerissa*, suggesting that a large portion of the butterfly groups comprised of rare species. The solid line on the plot represents the observed data, revealing more rare species than might be expected under an evenly distributed community. This pattern suggests low species evenness, as a few species are disproportionately abundant, while the majority are rare. Despite this unevenness, the high number of species indicated that the community still exhibited substantial species richness. Essentially, the community was diverse in terms of the number of species, but the dominance of a few species reduced the overall balance of species distribution.

**Table 2:** The diversity indices calculation of 5 different butterfly families.

| Family | H' | D | Dmg | J' |
| --- | --- | --- | --- | --- |
| Pieridae | 1.398 | 0.611 | 1.309 | 0.607 |
| Lycaenidae | 0.833 | 0.419 | 0.76 | 0.465 |
| Nymphalidae | 1.759 | 0.707 | 2.831 | 0.634 |
| Papilionidae | 0.295 | 0.118 | 0.553 | 0.213 |
| Hesperiidae | 1.099 | 0.667 | 1.82 | 1 |
| Total | 2.317 | 0.839 | 4.962 | 0.633 |

**Figure 8:**
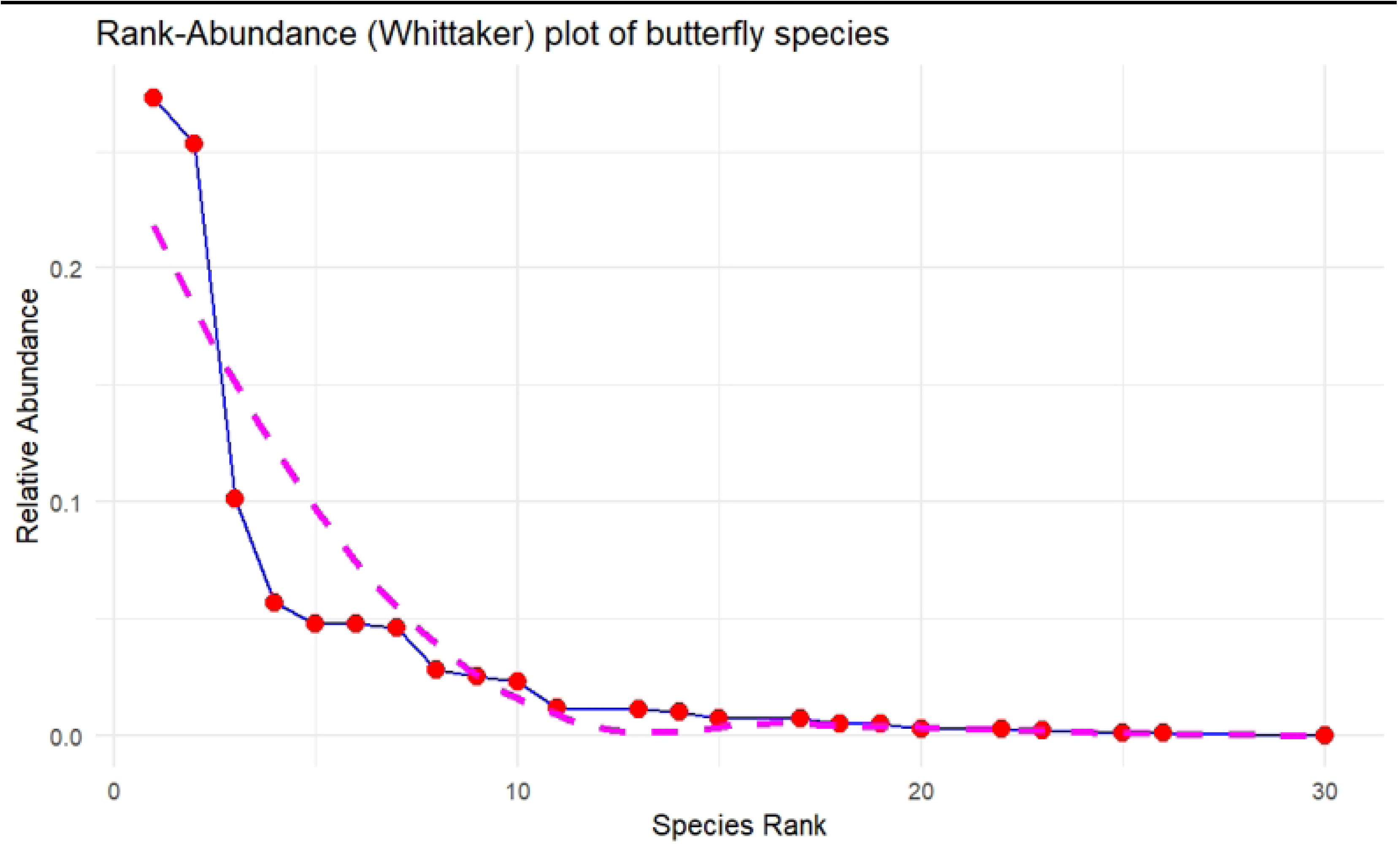
Rank-abundance plot of butterfly species observed.

Nymphalidae family was the most diverse, with the highest Shannon-Wiener index (H’ = 1.759), richness (Dmg = 2.831), and evenness (J’ = 0.634) among 16 species. Pieridae also showed high diversity (H’ = 1.398), richness (Dmg = 1.309), and evenness (J’ = 0.607) across 10 species. Hesperiidae, with only 3 species, had evenness (J’ = 1), while Lycaenidae and Papilionidae showed lower diversity (H’ = 0.833 and 0.295) and richness (Dmg = 0.76 and 0.553). Papilionidae had the lowest evenness (J’ = 0.213) with 4 species only. Overall, the butterfly community was fairly diverse (H’ = 2.317) with as many as 39 species recorded.

Grassland had the highest species richness, led by Pieridae and Lycaenidae followed by Shrubland and forest being moderately rich, with mostly Nymphalidae and Pieridae contributing (Figure 9). Agricultural land had lower richness, dominated by Pieridae, while wetlands had the fewest species. Pieridae dominated all habitats except shrubland.

**Table 3:** The Shannon Diversity Index (H’) and Pielou’s Evenness Index of the overall butterfly richness in 5 different habitats.

| Habitat | Shannon Diversity Index (H') | Pielou's Evenness Index (J) |
| --- | --- | --- |
| Agricultural land | 2.0628 | 0.7281 |
| Shrubland | 2.0263 | 0.7010 |
| Grassland | 1.1779 | 0.4463 |
| Forest | 1.7242 | 0.6367 |
| Wetland | 1.4195 | 0.6827 |

**Figure 9:**
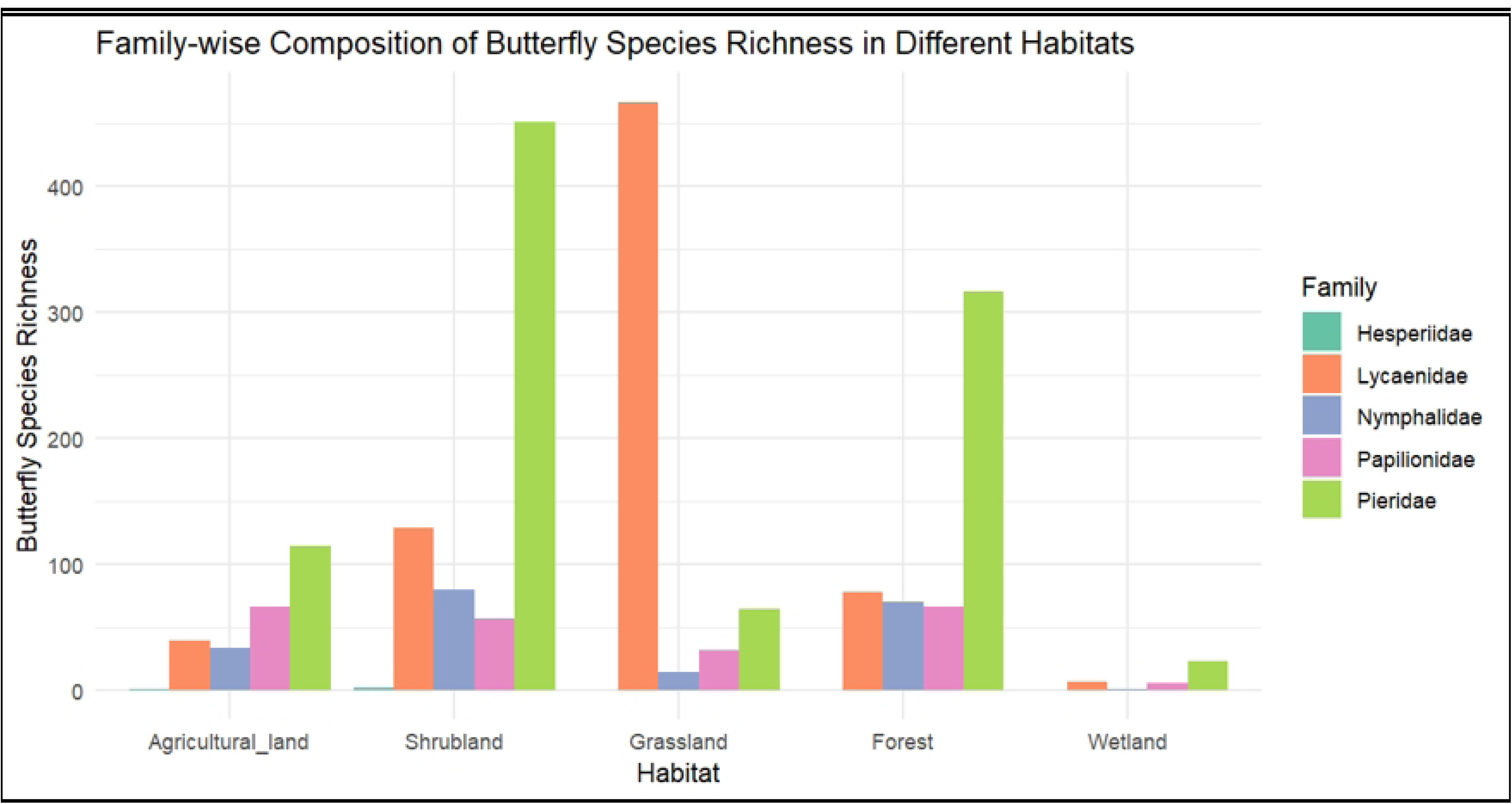
The family-wise categorization of butterfly richness in 5 different habitats.

In terms of habitat, agricultural land showed the highest diversity (H’ = 2.0628) and evenness (J = 0.7281), indicating a well-balanced species distribution. Shrubland followed closely with similar diversity (H’ = 2.0263) and slightly lower evenness (J = 0.701). Forest habitat displayed moderate diversity (H’ = 1.7242) and evenness (J = 0.6367). Wetlands had fewer species but showed a balanced composition with high evenness (J = 0.6827) and lower diversity (H’ = 1.4195). Grasslands were the least diverse (H’ = 1.1779) and even (J = 0.4463), with only a few species dominating.

Butterfly species were observed in different plant types—native, alien, planted, invasive, and weeds—and these had fundamental effects on their distribution. Native plants like *Clerodendrum viscosum* and *Dendrocalamus* attracted species such as *Necrophora chinense* and *Pachliopta aristolochiae.* Invasive plants, including *Mikania micrantha*, were preferred by butterflies like *Papilio polytes* and *Hypolimnas bolina*. Planted species, such as *Citrus limon*, attracted butterflies like *Gonepteryx rhamni*, while weeds like *Oxalis corniculata* were favored by species like *Pieris brassicae*. These showing a complex but intimate assocations between plant types and butterfly species.

## Discussion

Considering the time of the year and the limited timeframe for the study, finding of 30 moth species from six families, covering 26 genera, and 39 butterfly species from five families, spanning 31 genera, highlights a very rich Lepidoptera diversity in Lumbini. This diversity represents 5.88% of butterflies and 0.5% of moths recorded in Nepal.

### Dominance and Diversity of Moth Species

The Eribidae and Noctuidae families dominance covering 70% of all families recorded (Figure 5), show consistency with previous studies, such as those by Sondhi *et al.* [56], at Shendurney and Ponmudi in Kerala, India’s Agastyamalai Biosphere Reserve, where Eribidae was also identified as the most abundant moth family. Similarly, Ahmed *et al.* [1] reported that Eribidae was among the most commonly recorded moth families in the Mahamaya Reserve Forest in Assam. Following Eribidae, the Noctuidae family accounted for 29.79% of the moth species, with 8 different species recorded. Noctuidae is less diverse than Eribidae but makes a major contribution to the community, as evidenced by its moderate diversity. Other recorded families include Geometridae and Crambidae, each representing 5.32% of the total. Although these families have fewer species, they are evenly spread, as shown by their lower diversity but high evenness. Pyralidae, with 3.19% of the population and two species, show the lowest diversity (H’ = 0.64) but the highest evenness (J’ = 0.92), indicating a balanced but limited representation.

Some families, such as the Erebidae and Noctuidae, appear to be more adapted to the environment of the lemon farm and its surrounding plants, as evidenced by their dominance in the recordings (Appendix 2). The lemon farm, characterized by a semi-urban agricultural land/grassland with excessive growth of native/alien vegetation like *Imperata cylindrica* and *Mikania micrantha*, moth species may have adjusted to the altered plant ecology—doing well in disturbed or semi-natural habitats [35]. Likewise, several moths may choose certain habitats and food supplies provided by the natural grass species *Imperata cylindrica*. Consequently, the moth species that have been observed most probably are generalists, which can survive in disturbed habitats like farmland, adapting to the specific flora that is present [26].

Overall, the moth community has shown huge diversity with 30 species’ assemblage despite varying contributions from families. *Spilosoma strigatula* (25 individuals) and *Sesamia inferens* (18 individuals) are the most abundant species of this assemblage. The dominance of *S*. *strigatula* aligns with the studies by Sanyal *et al.* [46], who noted it as a major agricultural pest and a detector species rather than an indicator species.

The presence of the moth species that have been seen may have been impacted not only by the environment but also by varying weather patterns. During the full moon period (24 February 2024), for instance, only three species on average are recorded. For the night of the full moon, when the trap was examined the next day only one species was found! Whereas for other nights on average, six species have been recorded. According to Williams and Singh [66], when strong lights are used to attract insects at night, the number of species captured is substantially more during the new moon compared to the full moon. This is because when there is no moonlight during a new moon, artificial light is more appealing to insects, but when there is a full moon, artificial light is less effective and attracts comparatively fewer insects. Similarly, a complete cloud cover on 27 February 2024, in contrast to a full moon, resulted in a larger diversity of moths being reported, with seven distinct species being identified. Watkins [65] found a positive correlation between more cloud cover and high moth activity. This is probably because clouds may cover natural light sources like moonlight, which makes the light traps stand out and attracts moths looking for a dependable source of light during the dark [38]. Also cloud cover may help trap the heat of the day making nights warmer, therefore increased moth movement.

A few Fairly Common (FC) species, like *Spilosoma strigatula* and *Sesamia inferens*, make up a significant portion of the total, indicating their role as well-established pests with important ecological impacts. In contrast, most species are classified as Very Rare (VR) but contribute far less to the overall population, likely due to their restricted distribution and specialized habitat needs [32]. FC species, with their broader ecological niches and adaptability, have a more prominent presence in the ecosystem [32]. VR species like *Cretonotos transiens* often have specialized needs, resulting in limited distributions and smaller populations, highlighting their role in preserving genetic and ecological diversity but also their vulnerability to environmental changes [59]. Rare (R) species, such as those from the *Asura-miltochrista* and *Eressa confinis*, have moderate recordings, indicating they are less common than FC species but still play a significant role in the environment. While a few common species dominate, the presence of many rarer species adds ecological complexity, emphasizing the importance of habitat preservation for sustaining biodiversity.

Several environmental factors likely contribute to the rank abundance pattern observed (Figure 6), where a few dominant moth species are much more common than others. Environmental conditions, such as temperature, humidity, and vegetation, may favor certain species, leading to their dominance in the study area [22]. Dominant species may have a competitive edge due to life history traits such as higher reproduction rates, broader feeding preferences, or better adaptation to the local environment [52]. Additionally, abundance shifts could be influenced by certain species being more attracted to the light trap due to their nocturnal behavior or increased sensitivity to light [62]. Species near the lower end of the rank-abundance curve may be rarer or more specialized, accounting for their lower presence in the sample [8]. The unequal distribution of abundances in the plot likely results from environmental suitability, competitive advantages, and potential sampling bias.

### Dominance and Diversity of Butterfly Species

The Nymphalidae and Pieridae family combined cover 67% of the total recorded species. The data reveal a diverse butterfly community, with common species like *Pieris brassicae* thriving very well and species like *Junonia almana* and *Eurema hecabe* showing limited distribution as VR species. A similar diversity was observed in Thakurbaba Municipality, Bardiya, conducted in similar habitats in the lowlands of western Nepal, where Nymphalidae was also dominant [40]. Similarly, Sharma and Paudel [48] found that Nymphalidae as the dominant family in Kumakh Rural Municipality, Salyan District, Nepal. Despite the varied geography and climate in the Manang Region, Shrestha *et al.* [50] also observed Nymphalidae as the most species-rich family. This is also very interesting because of similar results despite of great altitudinal differences. This dominance is likely due to their adaptability to various habitats, diverse larvae host plants, and high reproductive rates, allowing them to thrive in both tropical and temperate regions [29].

The Rank-Abundance plot of butterfly species reveals a community where a few species dominate with high relative abundances such as the *Pieris canidia* and *Zizina otis*, while many others are rare. This pattern suggests a diverse ecosystem, where dominant species likely thrive due to better adaptation to environmental conditions or greater resilience to change [5]. The steep decline in relative abundance followed by a gradual slope highlights the presence of both common and rare species, highlighting the ecosystem’s complexity and richness, and emphasizing the need to protect these rare species to maintain biodiversity and resilience. *Pieris canidia* (C) was the most abundant butterfly species, comprising 27.29% of all individuals, closely followed by *Zizina otis* (C) at 25.31% (Figure 4). Shrestha *et al* [49] also identified *Pieris canidia* as the most abundant species, with *Aglais cashmerensis* as the second in different sacred forests of Kathmandu valley. However, *Aglais cashmerensis* was not recorded in this study, likely because it was still hibernating. According to Riyaz and Sivasankaran [44], this species hibernates from December to February in sheltered locations, becoming active in March as temperatures rise, which may explain the timing of the current study.

In this study, grassland exhibits the highest species richness, particularly in the Lycaenidae and Pieridae families (Figure 6), while agricultural land and shrubland demonstrate the greatest species diversity compared to grasslands. Agricultural land and wetlands show the highest species evenness at 0.7281 and 0.6827 respectively, while grassland has the lowest at 0.4463. The abundance of Lycaenidae and Pieridae in grassland and shrubland is tied to their unique ecological conditions. Open habitats like grassland offer ample nectar and host plants, providing the sunny environments essential for Pieridae butterflies like *Eurema brigitta* and *Leptosia nina*, which rely on plants such as *Oxalis corniculata* and *Lippia alba* (Table 4). Similarly, Lycaenidae species like *Zizina otis* and *Castalius rosimon* thrive in shrublands, benefiting from diverse plant life, including *Acmella uliginosa* and *Rungia pectinata*. These habitats support various microhabitats, allowing diverse species within these families to coexist and flourish, contributing to the observed high species richness [2]. The high species diversity in agricultural land and shrubland reflects a well-balanced community structure with many species distributed evenly, promoting ecological stability and biodiversity [58]. This distribution of butterfly species across habitats highlights the significant role of plant communities with different phenophases influencing butterfly community structure and abundance.

**Table 4:**
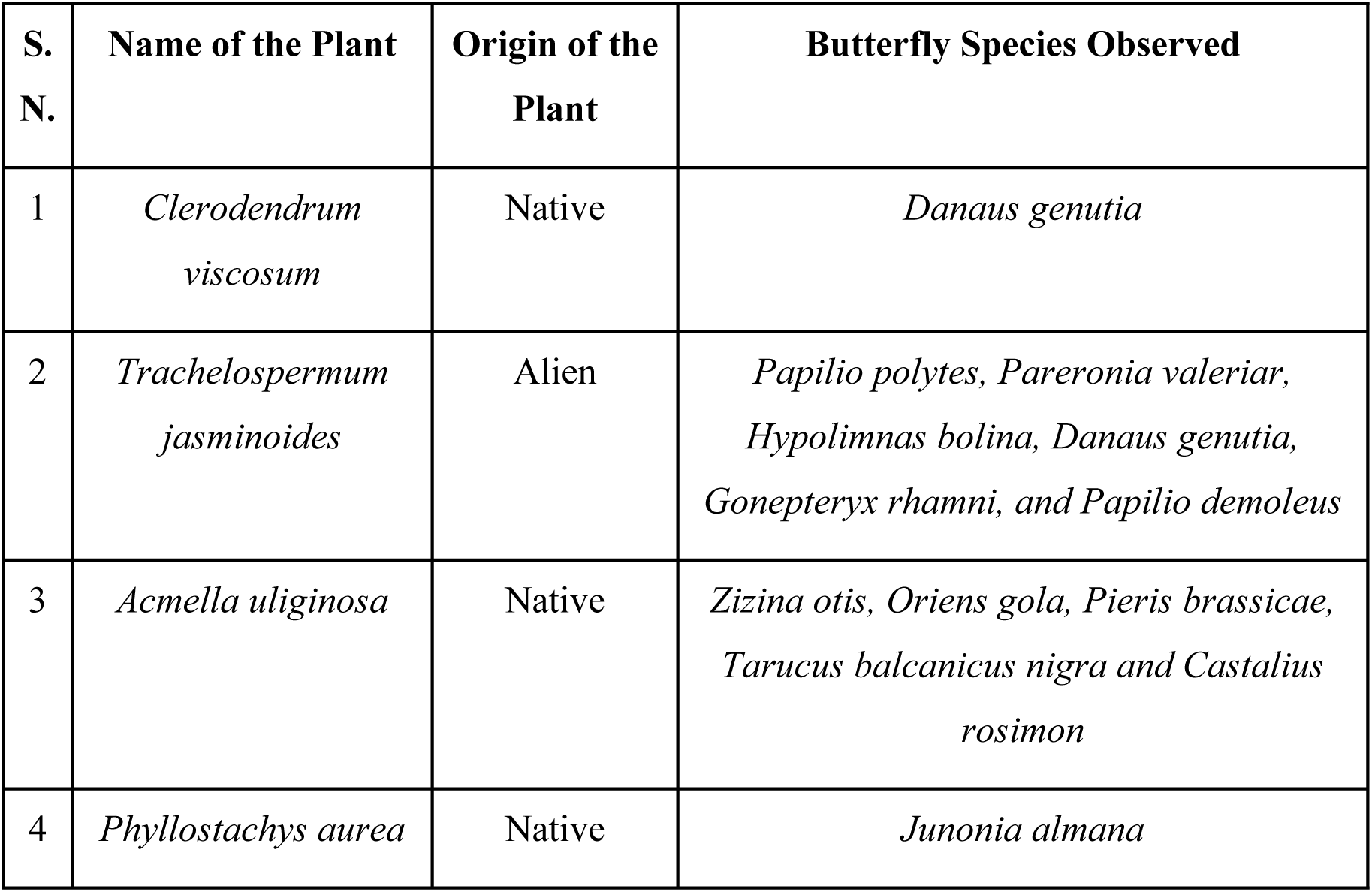
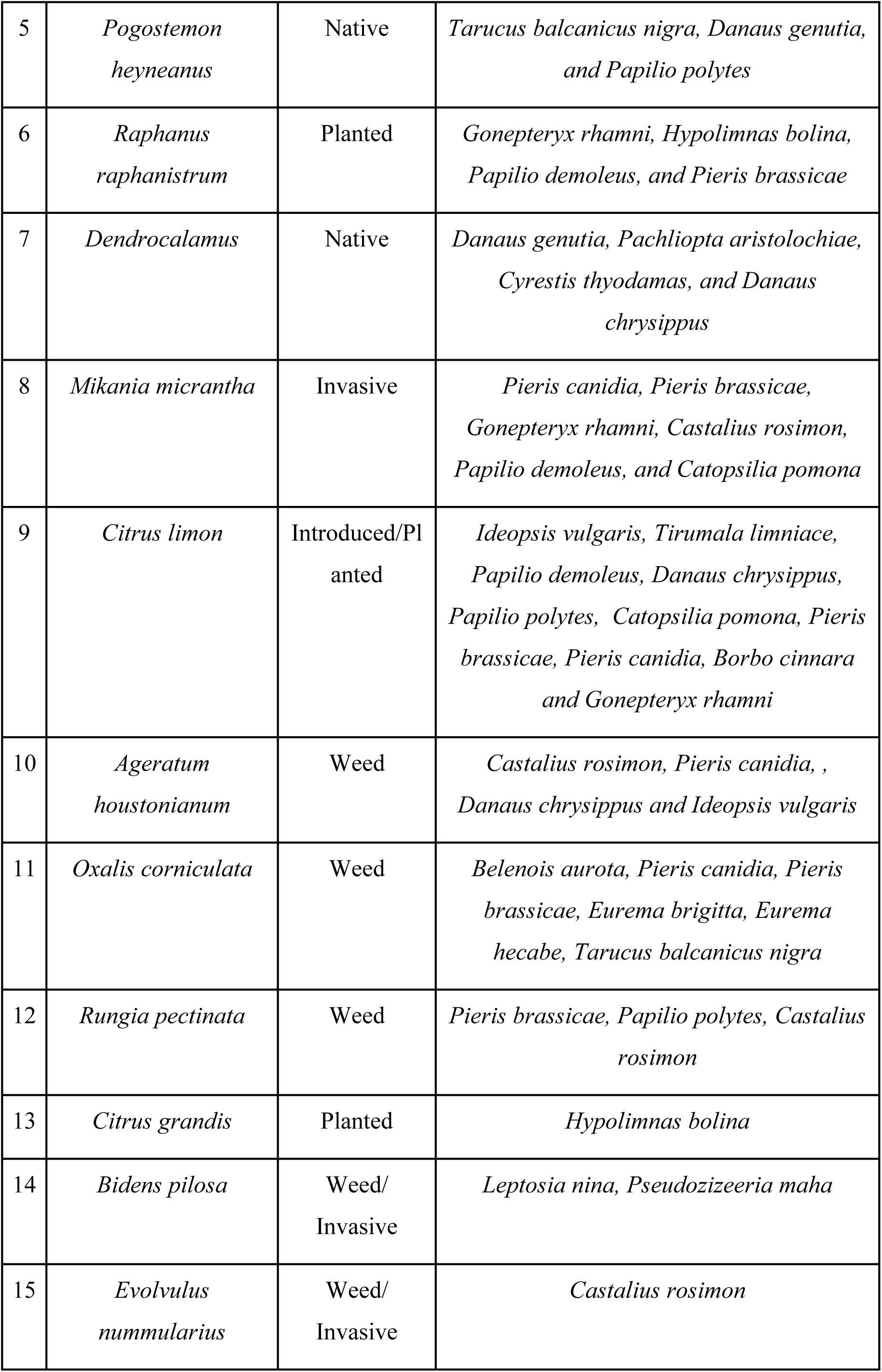
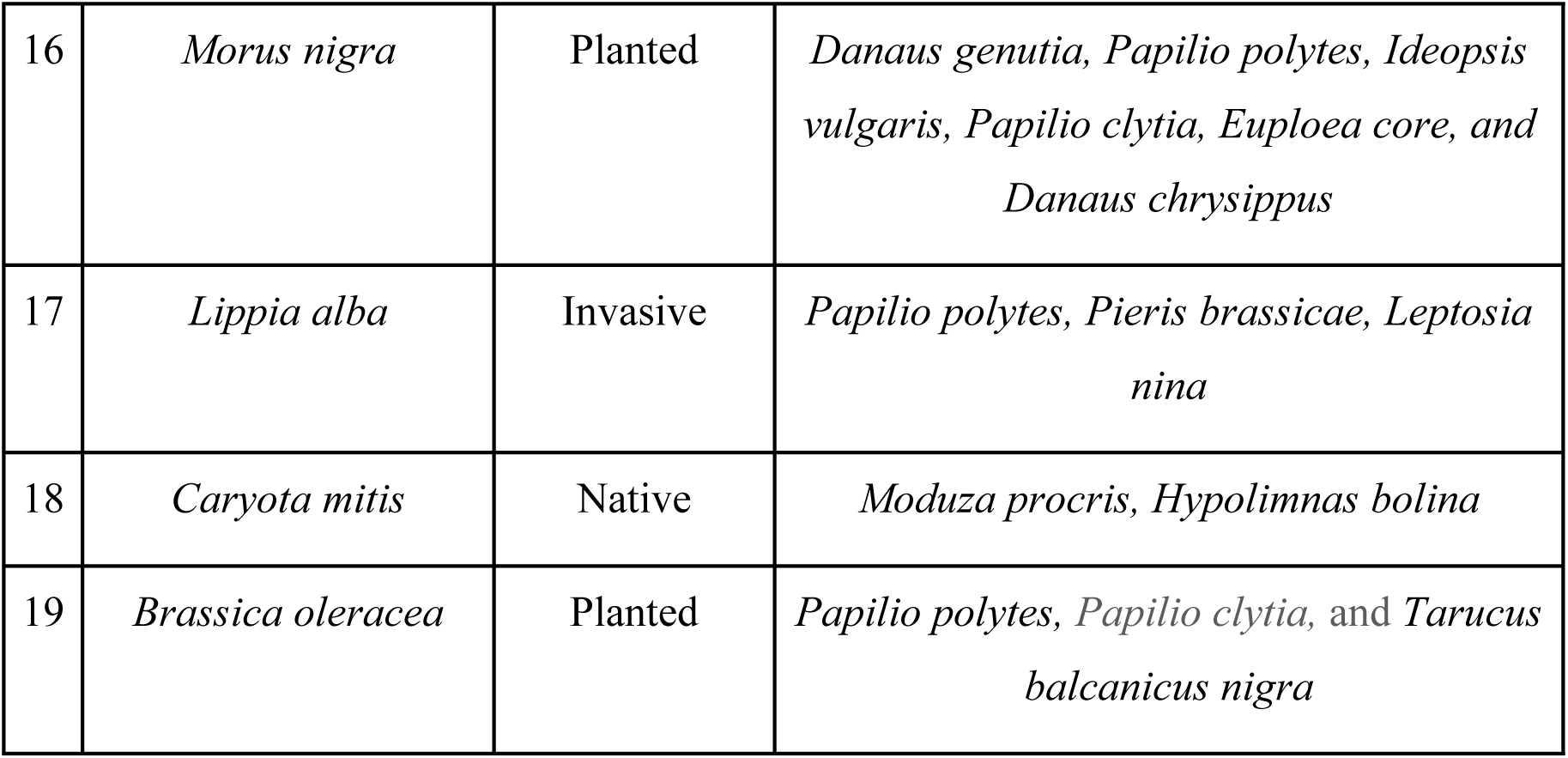
The total vegetation recorded with their origin and the butterfly species that visited the most.

Agricultural and shrub areas, with diverse plant species such as *Raphanus raphanistrum, Citrus limon, Citrus grandis, Morus nigra, Clerodendrum viscosum, Trachelospermum jasminoides, Pogostemon heyneanus,* and *Ageratum houstonianum*, offer abundant nectar and food sources. This plant diversity promotes a high abundance and richness of butterflies, highlighting the importance of plant diversity for supporting healthy butterfly populations [57]. In a study carried out in Nagpur, Central India, Tiple and Khurad [61] showed that agricultural land supports a wider variety of species compared to grasslands similar to this study. However, in contrast, studies by Paudel, Shah, and Bashyal [43] found that forest habitats had the highest species diversity, while farmland had the lowest. The current research may differ because the surveyed agricultural area was devoid of harmful pesticides, possibly attracting a wider variety of species. The forest covered only a small portion of our study area, therefore less diversity of butterflies. *Pieris canidia*, *Papilio polytes*, *Pieris brassicae*, and *Castalius rosimon* were the most frequent species across habitats, likely due to their adaptation to semi-urban areas, the abundance of suitable host plants from families like Brassicaceae and Rutaceae, and their rapid reproductive rates [30]. Their diverse feeding habits on readily available nectar sources also support their survival. According to Priyadarshana *et al.* [43], *Pieris canidia* and *Pieris brassicae* thrive in farmlands as important pests of cruciferous crops. Additionally, field edges and blooming weeds provide extra nutrients and habitat, further increasing their abundance [11].

The findings in Lumbini highlight the intricate relationships between Lepidoptera species and their surrounding habitats, reinforcing the vital role that plant diversity and environmental conditions play in shaping their populations. The dominance of the Nymphalidae family, commonly seen across Nepal, along with the rich diversity of moths in Lumbini’s farmlands, illustrates how habitat variation and management practices significantly influence species composition. The presence of diverse plant species supports a wider range of butterflies and moths, while environmental conditions such as temperature, moisture levels, and land use patterns further dictate which species thrive in particular habitats.

The adaptability of moth families such as Erebidae and Noctuidae, especially in disturbed environments, highlights their resilience to environmental changes. These families can persist in habitats altered by human activity or natural disturbances, which indicates their ecological flexibility and ability to thrive in a variety of niches. This adaptability not only highlights their capacity to survive in less-than-ideal conditions but also reveals how some species may even benefit from certain disturbances, allowing them to become more widespread.

Overall, these observations emphasize the importance of maintaining diverse and well-managed habitats to support a wide range of Lepidoptera species. By understanding the specific ecological needs of butterflies and moths, conservation efforts can be more effectively tailored to preserve both common and rare species. The findings also suggest that while some species show resilience to changing environments, others may be more vulnerable, making habitat preservation and sustainable land management crucial for maintaining biodiversity

## Conclusion

The study revealed a rich and diverse assemblage of Lepidoptera species across various habitats in Lumbini, highlighting significant differences in family representation, species abundance, and diversity. Among moths, the Erebidae family was the most abundant, followed by Noctuidae, while butterflies were dominated by the Pieridae and Lycaenidae families, particularly in grassland and shrubland habitats. These patterns highlights the distinct ecological roles and behaviors of moths and butterflies, which shape their species richness, evenness, and overall diversity.

Butterflies, being diurnal and often more specialized, showed lower species richness and evenness, with certain species dominating due to the availability of host plants and seasonal variations. In contrast, moths, being primarily nocturnal and more generalist, exhibited higher species richness and evenness, occupying a broader range of ecological niches and demonstrating greater adaptability to seasonal and environmental changes.

The impact of habitat types—such as agricultural lands, shrublands, forests, wetlands, and grasslands—on butterfly and moth distributions was evident, with butterflies thriving more in grassland and shrubland areas, while moths displayed a more balanced distribution across diverse habitats. The study’s diversity indices reflect variations across these ecosystems, with some habitats supporting more balanced species distributions, thereby highlighting the importance of habitat in determining Lepidoptera diversity and abundance. Additionally, the relationship between environmental factors, such as seasonal weather conditions and moon phases, and moth diversity, richness, and evenness was explored. Moths demonstrated a stronger adaptability to these factors, which shaped their ecological preferences and allowed them to occupy diverse habitats.

The findings also highlight the critical link between plants and butterfly species, illustrating how vegetation and specific habitats influence species diversity and distribution. This study enhances the understanding of the region’s biodiversity and ecological dynamics, particularly regarding how different ecosystems and environmental conditions structure both moth and butterfly communities. The observed species fluctuations highlight the need for ongoing monitoring to better grasp the ecological processes that affect these intricate populations.

## Recommendations

1. Extending the study period to cover a broader range of seasons would allow for a more precise understanding of Lepidoptera diversity, particularly for butterflies, whose populations are heavily influenced by seasonal fluctuations. This expanded timeframe would capture the full spectrum of their abundance, providing a more comprehensive dataset. Similarly, conducting detailed habitat studies on moths can yield critical insights into their specific ecological preferences and behavior. Regular multi-seasonal monitoring is essential for tracking shifts in species richness and evenness over time, offering a clearer picture of how habitat dynamics influence Lepidoptera populations.
2. To sustain biodiversity and ensure the survival of both moth and butterfly species, it is crucial to integrate habitat conservation with sustainable land-use practices. Protecting diverse habitats such as shrublands, wetlands, forests, and flowering areas will attract a broader range of species. Recommendations for specific plants that attract butterflies and moths can be made for public gardens, contributing to urban biodiversity and ecosystem health.
3. Improving species identification techniques through advanced categorization methods will also reduce errors, ensuring more accurate data on moth diversity. Accurate identification is vital for understanding species distribution patterns and informing conservation strategies.
4. Educational efforts play a pivotal role in fostering long-term conservation awareness. Initiatives such as butterfly or moth houses, alongside educational programs in schools, can raise public awareness, support conservation efforts, and even promote ecotourism. These measures not only contribute to biodiversity preservation but also encourage community engagement and sustainable environmental practices.

## Acknowledgments

I thank NAMI College, University of Northampton for giving me this amazing opportunity to work on my project. I would also like to thank for the assistance of the staff at Lumbini Buddha Garden Resort for being helpful during the entirety of the fieldwork.

## Appendix 1 A Checklist of Moth Species and Their Abundance, Categorized By their Family, and Status Recorded

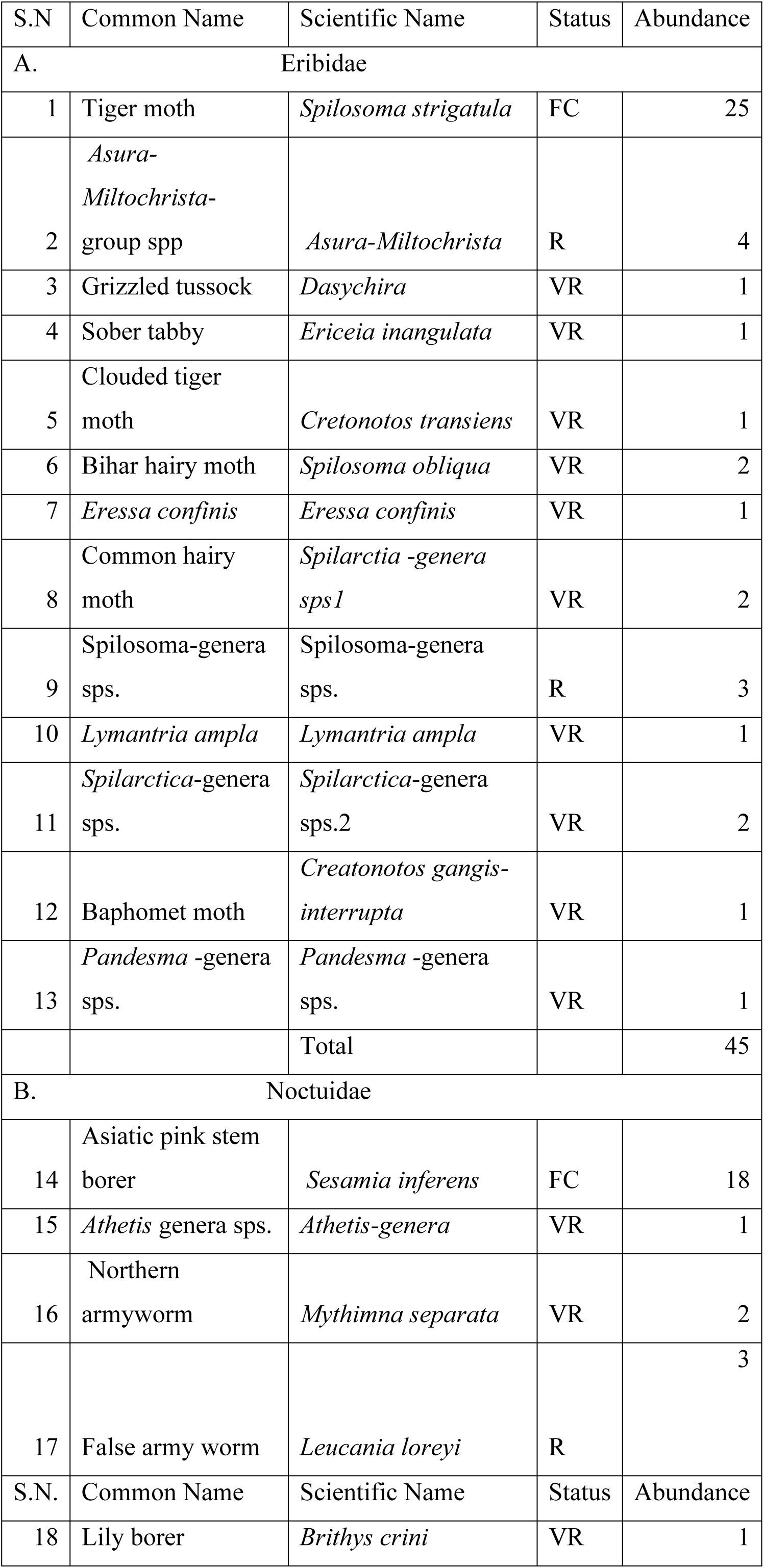
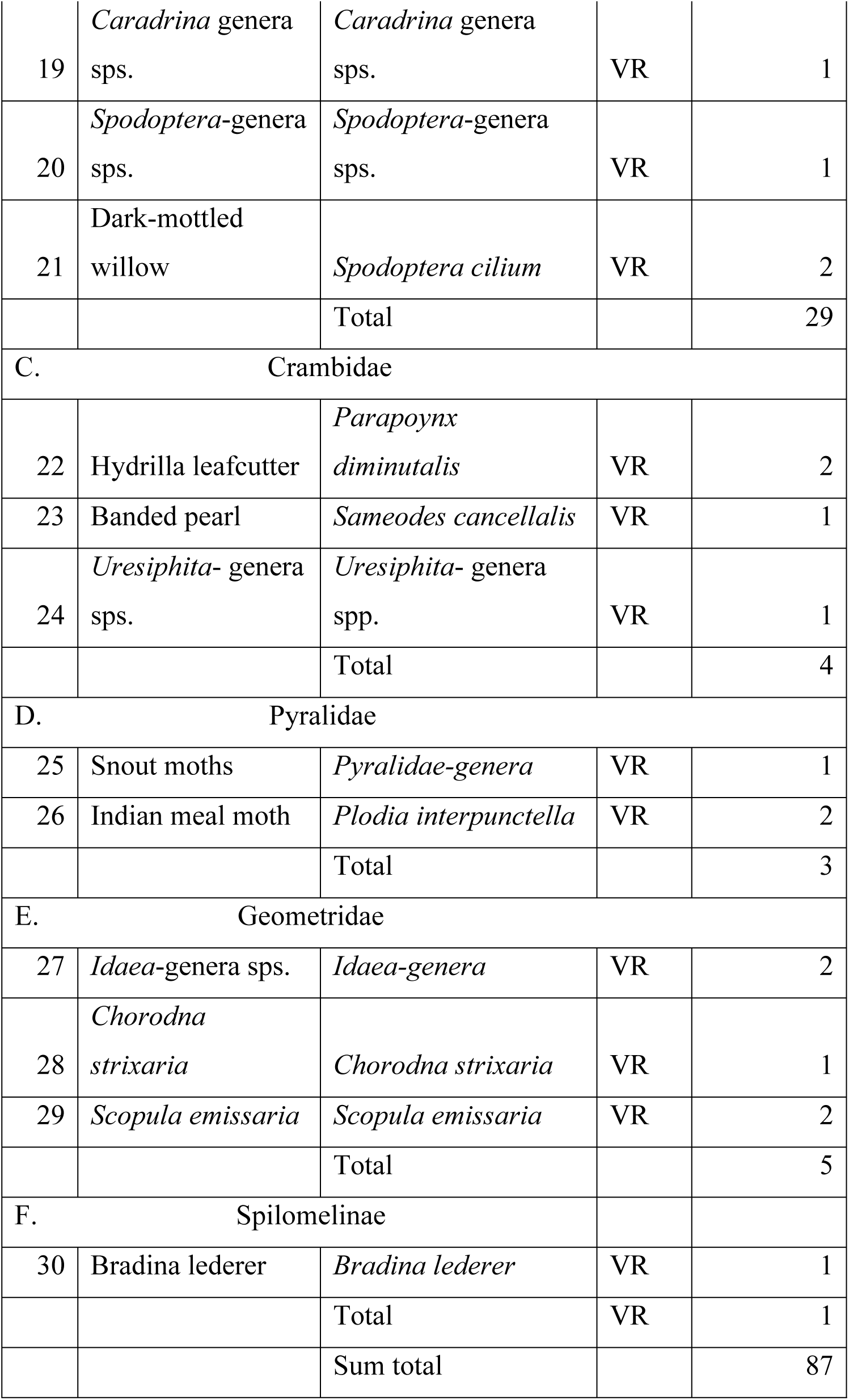

## Appendix 2 The List of the Butterfly Species Recorded Categorized in the Total Number of Families, and Individuals Found in Various Habitats such as; AgR=Agriculture land; ShR= Shrubland; GsR=Grassland; FsR= forest; WtL= Wetland; T=Total; ST= Sum Total

| S.N | Common Name | Scientific Name | Status | AgR | ShR | GsR | FsR | WtL | Total | Sum total |
| --- | --- | --- | --- | --- | --- | --- | --- | --- | --- | --- |
| A. | Pieridae |  |  |  |  |  |  |  |  |  |
| 1 | Common Gull | *Cepora nerissa* | FC | 7 | 8 | 0 | 8 | 0 | 23 | 969 |
| 2 | Indian Cabbage White | *Pieris brassicae* | C | 51 | 35 | 0 | 34 | 0 | 120 |  |
| 3 | Small Grass Yellow | *Eurema brigitta* | FC | 0 | 0 | 15 | 0 | 0 | 15 |  |
| 4 | Common Brimstone | *Gonepteryx rhamni* | C | 39 | 59 | 0 | 0 | 0 | 98 |  |
| 5 | Large Cabbage White | *Pieris canidia* | C | 0 | 280 | 0 | 275 | 23 | 578 |  |
| 6 | Common Wanderer | *Pareronia valeria* | C | 11 | 38 | 0 | 0 | 0 | 49 |  |
| 7 | Common Emigrant | *Catopsilia pomona* | FC | 6 | 9 | 0 | 0 | 0 | 15 |  |
| 8 | Caper White | *Pionner belenois aurota* | R | 0 | 3 | 7 | 0 | 0 | 10 |  |
| 9 | Wandering Psyche | *Leptosia nina* | *C* | 0 | 19 | 40 | 0 | 0 | 59 |  |
| 10 | Common Grass Yellow | *Eurema hecabe* | *VR* | 0 | 0 | 2 | 0 | 0 | 2 |  |
| B | Lycaenidae |  |  |  |  |  |  |  |  |  |
| 11 | Lesser Grass Blues | *Zizina otis* | *C* | 0 | 65 | 412 | 52 | 7 | 536 |  |
| 12 | Plain Cupid | *Luthrodes pandava* | *VR* | 0 | 0 | 2 | 0 | 0 | 2 |  |
| 13 | Common Pierrot | *Castalius rosimon* | *C* | 38 | 64 | 0 | 0 | 0 | 102 |  |
| 14 | Pea Blue | *Lampides boeticus* | *VR* | 1 | 0 | 0 | 0 | 0 | 1 |  |
| 15 | Pale Grass Blues | *Pseudozizeeria maha* | *FC* | 0 | 0 | 25 | 0 | 0 | 25 |  |
| 16 | Black-spotted Pierrot | *Tarucus balcanicus nigra* | *C* | 0 | 0 | 28 | 26 | 0 | 54 | 720 |
| C | Nymphalidae |  |  |  |  |  |  |  |  |  |
| 17 | Common Tiger | *Danaus genutia* | *C* | 25 | 43 | 0 | 33 | 0 | 101 |  |
| 18 | Great Eggfly | *Hypolimnas bolina* | *FC* | 3 | 0 | 0 | 22 | 0 | 25 |  |
| S.N | Common Name | Scientific Name | Status | Ag R | Sh R | Gs R | Fs R | Wt L | Total |  |
| 19 | Common Crow | *Euploea core* | *FC* | 0 | 21 | 0 | 0 | 0 | 21 |  |
| 20 | Blue Glassy Tiger | *Ideopsis vulgaris* | *R* | 3 | 4 | 0 | 0 | 0 | 7 |  |
| 21 | Commander | *Moduza procris* | *FC* | 0 | 5 | 0 | 9 | 0 | 14 |  |
| 22 | Common Leopard | *Phalanta phalantha* | *R* | 0 | 3 | 0 | 1 | 0 | 4 |  |
| 23 | Plain Tiger | *Danaus chrysippus* | *R* | 2 | 4 | 0 | 0 | 0 | 6 |  |
| 24 | Dark Evening Brown | *Melanitis phedima* | *VR* | 0 | 0 | 0 | 1 | 0 | 1 |  |
| 25 | Peacock Pansy | *Junonia almana* | *VR* | 0 | 0 | 0 | 2 | 0 | 2 |  |
| 26 | Grey Pansy | *Junonia atlites* | *VR* | 0 | 0 | 1 | 0 | 0 | 1 |  |
| 27 | Common Mapwing | *Cyrestis thyodamas* | *VR* | 0 | 0 | 0 | 2 | 0 | 2 |  |
| 28 | Bamboo Tree Brown | *Lethe europa* | *VR* | 0 | 0 | 0 | 0 | 1 | 1 |  |
| 29 | Common Evening Brown | *Melanitis leda* | *VR* | 0 | 0 | 1 | 0 | 0 | 1 |  |
| 30 | Great Evening Brown | *Melanitis zitenius* | *VR* | 0 | 0 | 1 | 0 | 0 | 1 |  |
| 31 | Oriental Blue Tiger | *Tirumala limniace* | *FC* | 0 | 0 | 11 | 0 | 0 | 11 | 199 |
| 32 | Plain Sailer | *Neptis cartica* | *VR* | 1 | 0 | 0 | 0 | 0 | 1 |  |
| D Papilionidae |  |  |  |  |  |  |  |  |  |  |
| 33 | Common Mormon | *Papilio polytes* | *C* | 64 | 54 | 30 | 61 | 4 | 213 | 227 |
| 34 | Common Mime | *Papilio clytia* | *R* | 1 | 3 | 0 | 0 | 0 | 4 |  |
| 35 | Common Rose | *Pachliopta aristolochiae* | *R* | 0 | 0 | 0 | 5 | 2 | 7 |  |
| 36 | Lime Swallowtail | *Papilio demoleus* | *R* | 1 | 0 | 2 | 0 | 0 | 3 |  |
| E Hesperidae |  |  |  |  |  |  |  |  |  |  |
| 37 | Indian Grizzled Skipper | *Spialia galba* | *VR* | 1 | 0 | 0 | 0 | 0 | 1 | 3 |
| 38 | Common Dartlet | *Oriens gola* | *VR* | 0 | 1 | 0 | 0 | 0 | 1 |  |
| 39 | Formosan Swift | *Borbo cinnara* | *VR* | 0 | 1 | 0 | 0 | 0 | 1 |  |
|  |  |  |  |  |  | Total abundance = |  |  |  | 2118 |

## Author contribution

Celeus Baral [Conceptualization (Lead), Data Curation (Lead), Formal Analysis (Lead), Methodology (Lead), Writing – Original draft (Lead)], Hem Sagar Baral (Writing – review and editing), Carol Inskipp (Writing – review and editing), Rajana Maharjan (Writing – review and editing).

